# CD47 as a potential biomarker for the early diagnosis of severe COVID-19

**DOI:** 10.1101/2021.03.01.433404

**Authors:** Katie-May McLaughlin, Denisa Bojkova, Marco Bechtel, Joshua D. Kandler, Philipp Reus, Trang Le, Julian U. G. Wagner, Sandra Ciesek, Mark N. Wass, Martin Michaelis, Jindrich Cinatl

## Abstract

The coronavirus SARS-CoV-2 is the cause of the ongoing COVID-19 pandemic. Most SARS-CoV-2 infections are mild or even asymptomatic. However, a small fraction of infected individuals develops severe, life-threatening disease, which is caused by an uncontrolled immune response resulting in hyperinflammation. Antiviral interventions are only effective prior to the onset of hyperinflammation. Hence, biomarkers are needed for the early identification and treatment of high-risk patients. Here, we show in a range of model systems and data from post mortem samples that SARS-CoV-2 infection results in increased levels of CD47, which is known to mediate immune escape in cancer and virus-infected cells. Systematic literature searches also indicated that known risk factors such as older age and diabetes are associated with increased CD47 levels. High CD47 levels contribute to vascular disease, vasoconstriction, and hypertension, conditions which may predispose SARS-CoV-2-infected individuals to COVID-19-related complications such as pulmonary hypertension, lung fibrosis, myocardial injury, stroke, and acute kidney injury. Hence, CD47 is a candidate biomarker for severe COVID-19. Further research will have to show whether CD47 is a reliable diagnostic marker for the early identification of COVID-19 patients requiring antiviral therapy.

## 1. Introduction

Severe acute respiratory syndrome coronavirus 2 (SARS-CoV-2) is causing the ongoing coronavirus disease 2019 (COVID-19) outbreak [Hokello et al., 2020; Chilamakuri & Agarwal, 2021], which has resulted in more than 100 million confirmed cases and more that 2 million confirmed COVID-19-associated deaths so far [Dong et al., 2020].

The first COVID-19 vaccines have been developed [Chilamakuri & Agarwal, 2021], and their roll-out has started in many countries. However, it will take a significant time until large parts of the world population will be vaccinated, and there is growing concern about the emergence of escape variants that can bypass immunity conferred by the current vaccines and previous SARS-CoV-2 infections [Andreano et al., 2020; Kemp et al., 2020; Liu et al., 2020; Weisblum et al., 2020; Sabino et al, 2021; Wibmer et al., 2021]. For the foreseeable future, there will thus be a need for improved COVID-19 therapies.

Currently, the therapeutic options for COVID-19 are still very limited [Chilamakuri & Agarwal, 2021; Rebold et al., 2021]. COVID-19 therapies can either directly inhibit SARS-CoV-2 replication or target other COVID-19-associated pathophysiological processes, such as corticosteroids that are anticipated to control COVID-19-related cytokine storm and hyperinflammation [Pum et al., 2021]. Dexamethasone and potentially other corticosteroids increase survival in patients who depend on oxygen support [RECOVERY Collaborative Group, 2020; WHO Rapid Evidence Appraisal for COVID-19 Therapies (REACT) Working Group, 2020]. In a controlled open-label trial, dexamethasone reduced mortality in patients receiving oxygen with (from 41.1% to 29.3%) or without (from 26.2% to 23.3%) mechanical ventilation, but increased mortality in patients not requiring oxygen support [RECOVERY Collaborative Group, 2020]. Other immunomodulatory therapy candidates are being tested, but conclusive results are pending [Rebold et al., 2021]. Further COVID-19 therapeutics under investigations include anticoagulants that target COVID-19-induced systemic coagulation and thrombosis (coagulopathy) [Hadid et al., 2020].

However, it would be much better to have effective antiviral treatments that reliably prevent COVID-19 disease progression to a stage when immunomodulators and anticoagulants are needed. The antiviral drug remdesivir was initially described to reduce recovery time from 15 to ten days and 29-day mortality from 15.2% to 11.4% [Beigel et al., 2020; Rebold et al., 2021]. However, other trials did not confirm this and conclusive evidence on the efficacy of remdesivir remains to be established [Rebold et al., 2021]. The JAK inhibitor baricitinib, which interferes with cytokine signalling, was reported to improve therapy outcomes in combination with remdesivir in a double-blind, randomised, placebo-controlled trial, in which patients were either treated with remdesivir plus baricitinib or remdesivir plus placebo [Kalil et al., 2020]. Moreover, convalescent sera and monoclonal antibodies are under clinical investigation for COVID-19 treatment [Tuccori et al., 2020; Devarasetti et al., 2021].

Ideally, antiviral therapies are used early in the disease course to prevent disease progression to the later immunopathology-driven stages [Weinreich et al., 2021]. However, only a small proportion of patients develops severe disease [Salzberger et al., 2020]. Therefore, biomarkers are required that identify patients, who will develop severe disease, as early as possible.

Here, we investigated the potential of the ubiquitously expressed cell surface glycoprotein CD47 as biomarker for the identification of high-risk COVID-19 patients. CD47 is the receptor of thrombospondin-1 (THBS1) and the counter-receptor for signal regulatory protein-α (SIRPα). CD47 interaction with SIRPα inhibits the activation of macrophages and dendritic cells and thrombospondin-1/ CD47 signalling inhibits T cell activation [Cham et al., 2020; Kaur et al., 2020]. High CD47 expression prevents immune recognition of cancer and virus-infected cells [Cham et al., 2020; Kaur et al., 2020].

## 2. Materials and Methods

### 2.1 Cell culture

Calu-3 cells (ATCC) were grown at 37°C in minimal essential medium (MEM) supplemented with 10% foetal bovine serum (FBS), 100 IU/mL penicillin, and 100 μg/mL of streptomycin. All culture reagents were purchased from Sigma. Cells were regularly authenticated by short tandem repeat (STR) analysis and tested for mycoplasma contamination.

Primary human bronchial epithelial cells were purchased from ScienceCell. For differentiation to air-liquid interface (ALI) cultures, the cells were thawed and passaged once in PneumaCult-Ex Medium (StemCell technologies) and then seeded on transwell inserts (12 well plate, Sarstedt) at 4×10^4^ cells/insert. Once cell layers reached confluency, medium on the apical side of the transwell was removed, and medium in the basal chamber was replaced with PneumaCult ALI Maintenance Medium (StemCell Technologies) including Antibiotic/Antimycotic solution (Sigma Aldrich) and MycoZap Plus PR (Lonza). During a period of four weeks, medium was changed and cell layers were washed with PBS every other day. Criteria for successful differentiation were the development of ciliated cells and ciliary movement, an increase in transepithelial electric resistance indicative of the formation of tight junctions, and mucus production.

### 2.2 Virus infection

SARS-CoV-2/7/Human/2020/Frankfurt (SARS-CoV-2/FFM7) was isolated and cultivated in Caco2 cells (DSMZ) as previously described [Hoehl et al., 2020; Toptan et al., 2020]. Virus titres were determined as TCID50/ml in confluent cells in 96-well microtiter plates [Cinatl et al., 2003; Cinatl et al., 2005].

### 2.3 Antiviral assay

Confluent layers of CaCo-2 cells in 96-well plates were infected with SARS-CoV–2 FFM7 at a MOI of 0.01. Virus was added simultaneously with tested compound. 24 h post infection, cells were fixed with acetone:methanol (40:60) solution and immunostaining of spike protein was performed to determine infection rate. Briefly, a monoclonal antibody directed against the spike protein of SARS-CoV-2 (1:1,500, Sino Biological) was detected with a peroxidase-conjugated anti-rabbit secondary antibody (1:1,000, Dianova), followed by addition of AEC substrate. The spike positive area was scanned and quantified by automated plate reader. The results are expressed as percentage of infection relative to virus control which received no drug.

### 2.4 Western blot

Cells were lysed using Triton-X-100 sample buffer, and proteins were separated by SDS-PAGE. Detection occurred by using specific antibodies against CD47 (1:100 dilution, CD47 Antibody, anti-human, Biotin, REAfinity™, # 130-101-343, Miltenyi Biotec), SARS-CoV-2 N (1:1000 dilution, SARS-CoV-2 Nucleocapsid Antibody, Rabbit MAb, #40143-R019, Sino Biological), and GAPDH (1:1000 dilution, Anti-G3PDH Human Polyclonal Antibody, #2275-PC-100, Trevigen). Protein bands were visualized and quantified by laser-induced fluorescence using infrared scanner for protein quantification (Odyssey, Li-Cor Biosciences).

### 2.5. qPCR

SARS-CoV-2 RNA from cell culture supernatant samples was isolated using AVL buffer and the QIAamp Viral RNA Kit (Qiagen) according to the manufacturer’s instructions. Absorbance-based quantification of the RNA yield was performed using the Genesys 10S UV-Vis Spectrophotometer (Thermo Scientific). RNA was subjected to OneStep qRT-PCR analysis using the Luna Universal One-Step RT-qPCR Kit (New England Biolabs) and a CFX96 Real-Time System, C1000 Touch Thermal Cycler. Primers were adapted from the WHO protocol29 targeting the open reading frame for RNA-dependent RNA polymerase (RdRp): RdRP_SARSr-F2 (GTG ARA TGG TCA TGT GTG GCG G) and RdRP_SARSr-R1 (CAR ATG TTA AAS ACA CTA TTA GCA TA) using 0.4 µM per reaction. Standard curves were created using plasmid DNA (pEX-A128-RdRP) harbouring the corresponding amplicon regions for RdRP target sequence according to GenBank Accession number NC_045512. For each condition three biological replicates were used. Mean and standard deviation were calculated for each group.

### 2.6 Data acquisition and analysis

Normalized protein abundance data from SARS-CoV-2-infected Caco-2 cells were derived from a recent publication [Bojkova et al., 2020]. Data were subsequently normalised using summed intensity normalization for sample loading, followed by internal reference scaling and Trimmed mean of M normalization. Mean protein abundance was plotted using the function *ggdotplot* of the R package ggpubr. P-values were determined by two-sided student’s t-test

Raw read counts from post-mortem samples of two COVID-19 patients and two healthy controls, as well as mock infected and SARS-CoV-2-infected Calu-3 cells, were derived from a recent publication [Blanco-Melo et al., 2020] via the gene expression omnibus (GEO) database (accession: GSE147507) and processed using DESeq2. Normalised gene counts were plotted using the function *ggdotplot* of the R package ggpubr. P-values were determined by two-sided student’s t-test

### 2.6 Literature review

Relevant articles were identified by using the search terms ‘CD47 aging’, ‘CD47 hypertension’, ‘CD47 diabetes’, and ‘CD47 obesity’ in PubMed (https://pubmed.ncbi.nlm.nih.gov). Original articles in English were included into the analysis, when they contained on the influence of aging, diabetes, diabetes, or obesity on CD47 expression levels and/ or the relevance of CD47 with regard to pathological conditions observed in severe COVID-19.

## 3. Results

### 3.1 SARS-CoV-2 infection results in enhanced CD47 expression

A publicly available proteomics dataset [Bojkova et al., 2020] indicated increased in CD47 expression in SARS-CoV-2-infected Caco2 colorectal carcinoma cells (Figure 1A). We also detected enhanced CD47 levels in SARS-CoV-2-infected primary human bronchial epithelial cells (HBEpiC) grown in air liquid interface (ALI) cultures [Bojkova et al., 2020A] and Calu-3 lung cancer cells (Figure 1B). Analysis of transcriptomics data from another study also indicated increased CD47 levels in SARS-CoV-2 -infected Calu-3 cells (Suppl. Figure 2) and in post mortem lung samples from COVID-19 patients (Figure 1C) [Blanco-Melo et al., 2020].

**Figure 1.**
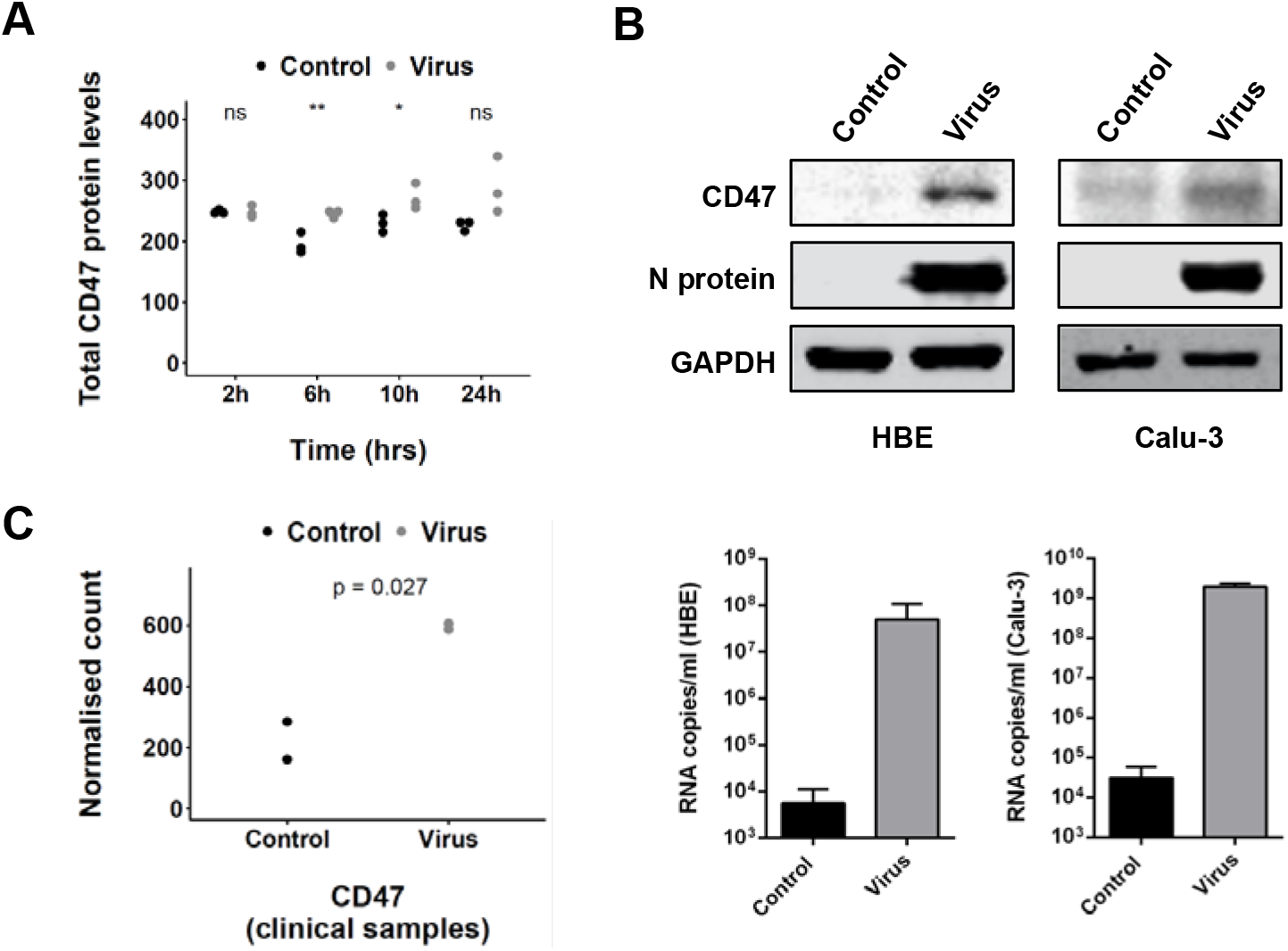
SARS-CoV-2 infection is associated with increased CD47 levels. A) TF protein abundance in uninfected (control) and SARS-CoV-2-infected (virus) Caco-2 cells (data derived from [Bojkova et al., 2020]. P-values were determined by two-sided Student’s t-test. B) CD47 and SARS-CoV-2 N protein levels and virus titres (genomic RNA determined by PCR) in SARS-CoV-2 strain FFM7 (MOI 1)-infected air-liquid interface cultures of primary human bronchial epithelial (HBE) cells and SARS-CoV-2 strain FFM7 (MOI 0.1)-infected Calu-3 cells. Uncropped blots are provided in Suppl. Figure 1. C) CD47 mRNA levels in post mortem samples from COVID-19 patients (data derived from [Blanco-Melo et al., 2020]). P-values were determined by two-sided Student’s t-test.

### 3.2 CD47 and COVID-19 risk factors

To further investigate a potential role of CD47 as biomarker indicating a high risk from COVID-19, we performed systematic literature searches on the relationship of CD47 and the known COVID-19 risk factors ‘ageing’, ‘diabetes’, and ‘obesity’.

#### 3.2.1 CD47 and ageing

The risk of severe COVID-19 disease and COVID-19 death increases with age [Hokello et al., 2020]. A literature search in PubMed (https://pubmed.ncbi.nlm.nih.gov, 17^th^ February 2020) using the terms ‘CD47’ and aging’ resulted in 62 hits (Suppl. Table 1). Eight of these articles contained information that support a link between age-related increased CD47 levels and an elevated risk of severe COVID-19 (Figure 2, Suppl. Table 1). One article suggested that alpha-tocopherol reduced age-associated streptococcus pneumoniae lung infection in mice by CD47 downregulation [Ghanem et al., 2015], which is in accordance with the known immunosuppressive functions of CD47 [Cham et al., 2020; Kaur et al., 2020].

**Figure 2.**
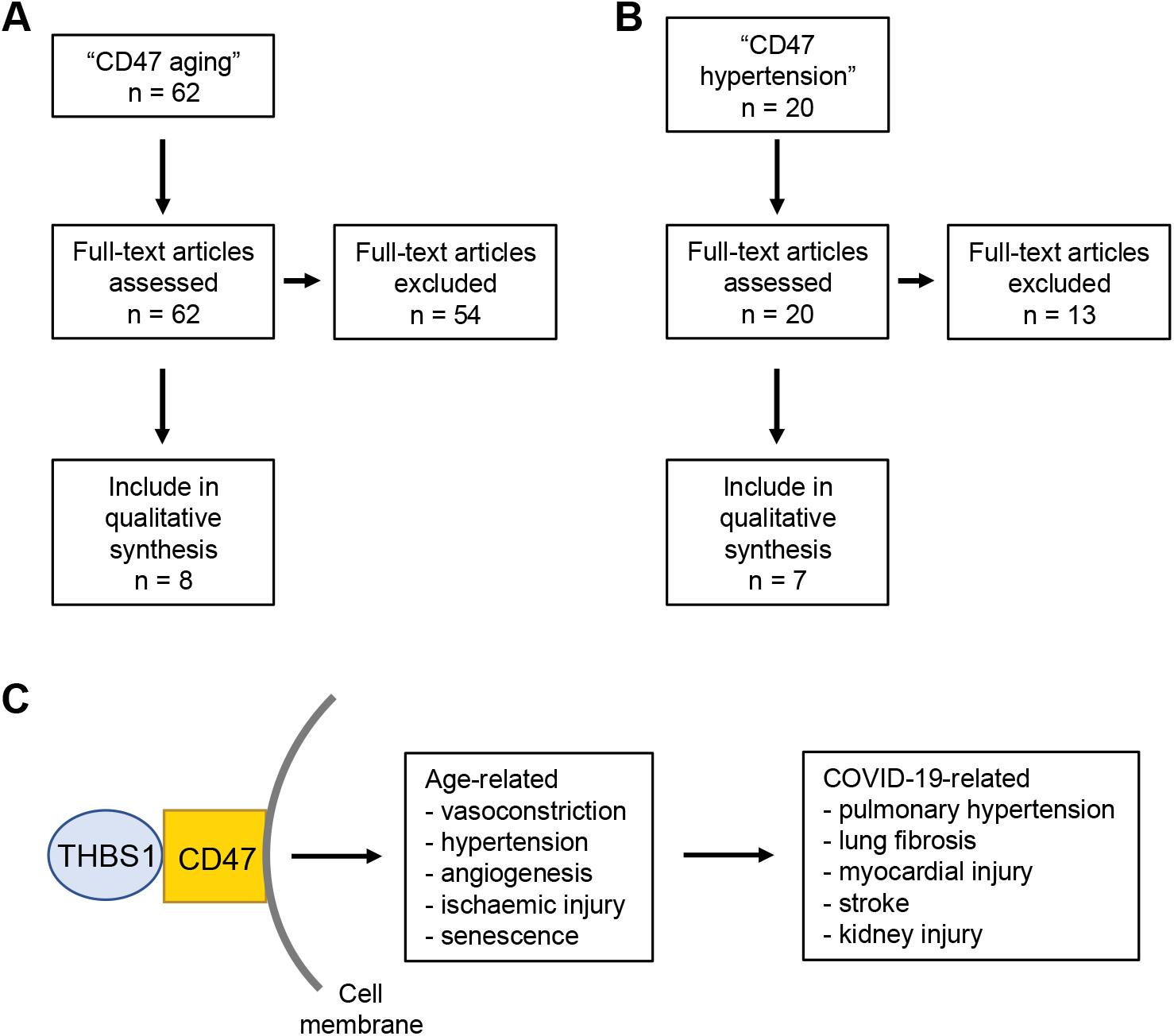
Results of the PubMed (https://pubmed.ncbi.nlm.nih.gov) literature search for “CD47 aging” (A) and “CD47 hypertension” (B). C) Overview figure of the data derived from the literature searches. Age-related increased CD47 levels may contribute to pathogenic conditions associated with severe COVID-19.

The remaining seven articles reported on age-related increased CD47 levels in vascular cells that are associated with reduced vasodilatation and blood flow (Suppl. Table 1), as CD47 signalling inhibits NO-mediated activation of soluble guanylate cyclase and in turn vasodilatation [Isenberg et al., 2008; Miller et al., 2010]. Since reduced vasodilatation can cause hypertension [Touyz et al., 2018], we performed a follow-up literature search using the search terms “CD47 hypertension” (Suppl. Table 2). This resulted in 20 hits, including a further seven relevant studies (Figure 2B, Suppl. Table 2).

Initial experiments showed that loss or inhibition of CD47 prevented age- and diet-induced vasculopathy and reduced damage caused by ischaemic injury in mice [Isenberg et al., 2007]. CD47-deficient mice indicated that CD47 functions as vasopressor and were also shown to be leaner and to display enhanced physical performance and a more efficient metabolism [Isenberg et al., 2009; Frazier et al., 2011]. In agreement, CD47 was upregulated in clinical pulmonary hypertension and contributed to pulmonary arterial vasculopathy and dysfunction in mouse models [Bauer et al., 2012; Rogers et al., 2017]. Age-related increased CD47 levels further affected peripheral blood flow and wound healing in mice [Rogers et al., 2013] and NO-mediated vasodilatation of coronary arterioles of rats [Nevitt et al., 2016]. Moreover, thrombospondin-1/ CD47 signalling was shown to induce ageing-associated senescence in endothelial cells [Gao et al., 2016; Meijles et al., 2017] and age-associated deterioration in angiogenesis, blood flow, and glucose homeostasis [Ghimire et al., 2020].

Increased CD47 levels were also detected in the lung of a sickle cell disease patient with pulmonary arterial hypertension, and vasculopathy and pulmonary hypertension were reduced in a CD47-null mouse model of sickle cell disease [Rogers et al., 2013a; Novelli et al., 2019]. Finally, anti-CD47 antibodies reversed fibrosis in various organs in mouse models [Wernig et al., 2017], which may be relevant in the context of COVID-19-associated pulmonary fibrosis [Leeming et al., 2021].

In addition to immunosuppressive activity, ageing-related increased CD47 levels may thus be involved in vascular disease, vasoconstriction, and hypertension and predispose COVID-19 patients to related pathologies such as pulmonary hypertension, lung fibrosis, myocardial injury, stroke, and acute kidney injury [Soto-Pantoja et al., 2013; Rogers et al., 2017a; Cruz Rodriguez et al., 2020; Fabrizi et al., 2020; Karmouty-Quintana et al., 2020; Scutelnic and Heldner, 2020; Leeming et al., 2021; Sanghvi et al., 2021; Shah et al., 2021].

#### 3.2.2 CD47 and diabetes

Diabetes has been associated with an increased risk of severe COVID-19 and COVID-19-related death [Shah et al., 2021]. A PubMed search for “CD47 diabetes” produced 47 hits, nine of which reported on increased CD47 levels in response to hyperglycaemia and/ or diabetes (Figure 3, Suppl. Table 3).

**Figure 3.**
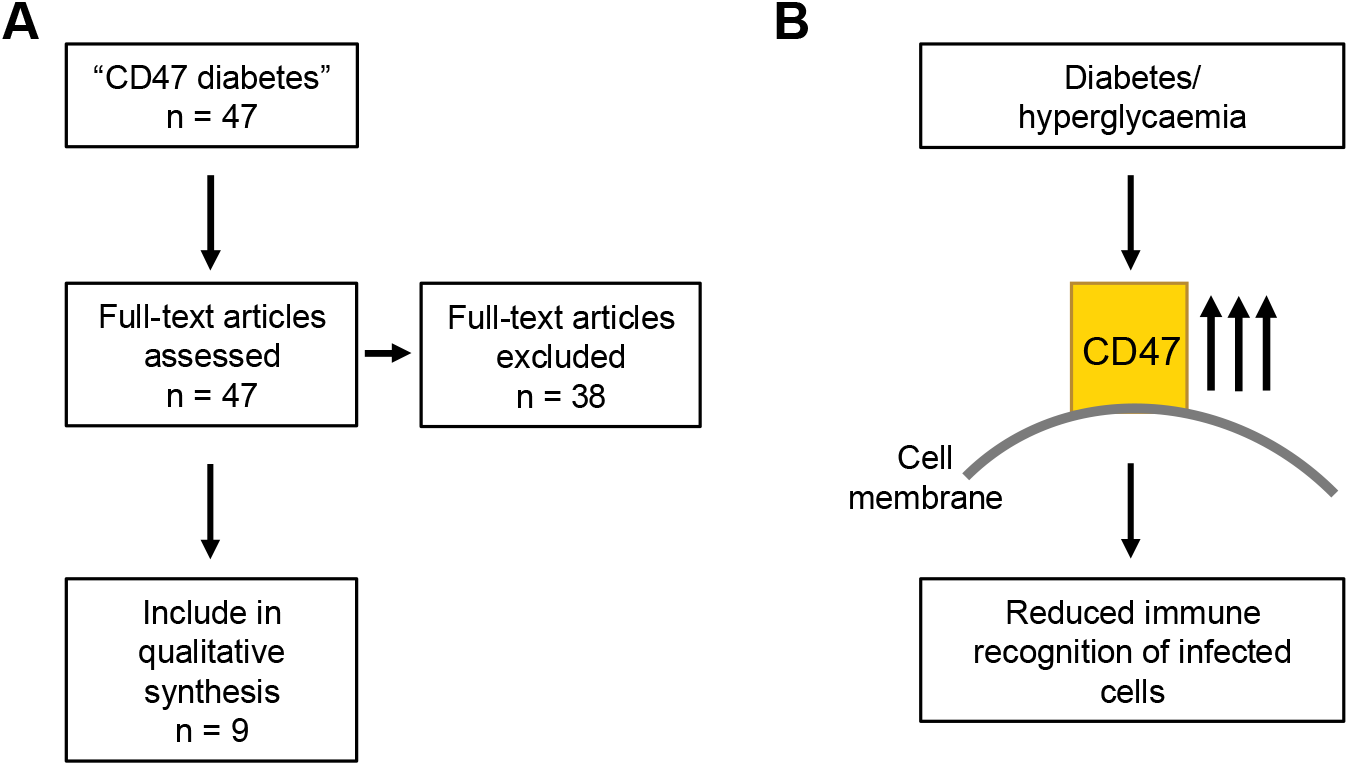
Results of the PubMed (https://pubmed.ncbi.nlm.nih.gov) literature search for “CD47 diabetes” (A). B) Overview figure of the data derived from the literature search. Hyperglycaemia- and diabetes-induced increased CD47 levels may contribute to immune escape of SARS-CoV-2-infected cells.

Hyperglycaemia protected CD47 from cleavage resulting in increased CD47 levels [Maile et al., 2008; 2009; 2010; 2012]. In agreement, increased CD47 levels were detected in various cell types and tissues in rat diabetes models and diabetes patients [Abdul-Rahman et al., 2012; Abu El-Asrar et al., 2013; Wang et al., 2014; Bitar, 2019]. Therefore, diabetes-induced increased CD47 levels may interfere with the recognition of SARS-CoV-2-infected cells by the immune system [Cham et al., 2020; Kaur et al., 2020].

#### 3.2.3 CD47 and obesity

As obesity is another risk factor for severe COVID-19 [Shah et al., 2021], we also performed a PubMed search for “CD47 obesity”, which resulted in eight hits, two of which provided potentially relevant information (Suppl. Table 4). Results indicated that CD47-deficient mice were leaner, probably as a consequence of elevated lipolysis [Maimaitiyiming et al., 2015; Norman-Burgdolf et al., 2020]. Hence, low CD47 levels may be associated both with lower weight and increased immune recognition of virus-infected cells [Maimaitiyiming et al., 2015; Cham et al., 2020; Kaur et al., 2020; Norman-Burgdolf et al., 2020], but there is no direct evidence suggesting that obesity may also directly increase CD47 expression. However, obesity may at least indirectly contribute to enhanced CD47 levels as risk factor for diabetes [Shah et al., 2021].

## 4. Discussion

Here, we show that SARS-CoV-2 infection is associated with increased CD47 expression in a range of model systems and in post mortem samples from COVID-19 patients. CD47 exerts immunosuppressive activity via interaction with SIRPα on immune cells and as thrombospondin-1 receptor [Cham et al., 2020; Kaur et al., 2020]. In this context, human CD47 expression is discussed as a strategy to enable the xenotransplantation of organs from pigs to humans [Cooper et al., 2019; Hosny et al., 2021]. Moreover, high CD47 expression is an immune escape mechanism observed on cancer cells, and anti-CD47 antibodies are under investigation as cancer immunotherapeutics [Feng et al., 2020; Kaur et al., 2020]. Due its immunosuppressive action, CD47 expression is also discussed as a target for the treatment of viral and bacterial pathogens including SARS-CoV-2 [Cham et al., 2020; Oronsky et al., 2020a; Tal et al., 2020]. Thus, our data indicating increased CD47 levels in a range of SARS-CoV-2 infection models and clinical samples further support the potential role of CD47 as drug target for the mediation of a more effective antiviral immune response.

Older age, diabetes, and obesity are known risk factors for COVID-19 morbidity and mortality [Hokello et al., 2020; Shah et al., 2021]. Hence, we performed a series of systematic reviews to identify potential connections between CD47 and these processes. Results indicated an ageing-related increase in CD47 expression, which may contribute the increased COVID-19 vulnerability in older patients [Hokello et al., 2020]. Moreover, high CD47 levels are known to be involved in vascular disease, vasoconstriction, and hypertension, which may predispose SARS-CoV-2-infected individuals to various conditions associated with severe COVID-19 related, including pulmonary hypertension, lung fibrosis, myocardial injury, stroke, and acute kidney injury [Soto-Pantoja et al., 2013; Rogers et al., 2017a; Cruz Rodriguez et al., 2020; Fabrizi et al., 2020; Karmouty-Quintana et al., 2020; Scutelnic and Heldner, 2020; Leeming et al., 2021; Sanghvi et al., 2021; Shah et al., 2021].

High CD47 levels have also been reported as a consequence of hyperglycaemia and diabetes, which may contribute to the high risk of severe COVID-19 in diabetic patients [Shah et al., 2021]. Although there is no known direct impact of obesity on CD47 levels, obesity is associated with an increased risk of diabetes and other ageing-related conditions such as hypertension that may result in elevated COVID-19 vulnerability [Shah et al., 2021].

## 5. Conclusions

Severe COVID-19 disease is the consequence of hyperinflammation (‘cytokine storm’) in response to SARS-CoV-2 infection [Jacques and Apedaile, 2020; Nowill and Campos-Lima, 2020; Gustine and Jones, 2021]. Hence, the optimal time window for antiviral intervention is as early as possible to prevent disease progression to severe stages driven by immunopathology [Weinreich et al., 2021]. Since the vast majority of cases are mild or even asymptomatic [Salzberger et al., 2020], biomarkers are required to identify and treat individuals early, who are at high risk of severe COVID-19.

Here, we have identified CD47 as a candidate biomarker for severe COVID-19. SARS-CoV-2 infection results in enhanced CD47 expression, which is known to interfere with the host immune response. Moreover, CD47 levels are elevated in groups at high risk from COVID-19 such as older individuals and individuals with hypertension and/ or diabetes. Thus, high CD47 levels may predispose these groups to severe COVID-19. Further research will have to show whether CD47 is a reliable diagnostic marker for the early identification of COVID-19 patients requiring antiviral therapy.

