## Supplementary material for "CD47 as a potential biomarker for the early diagnosis of severe COVID-19": Suppl Figure 1

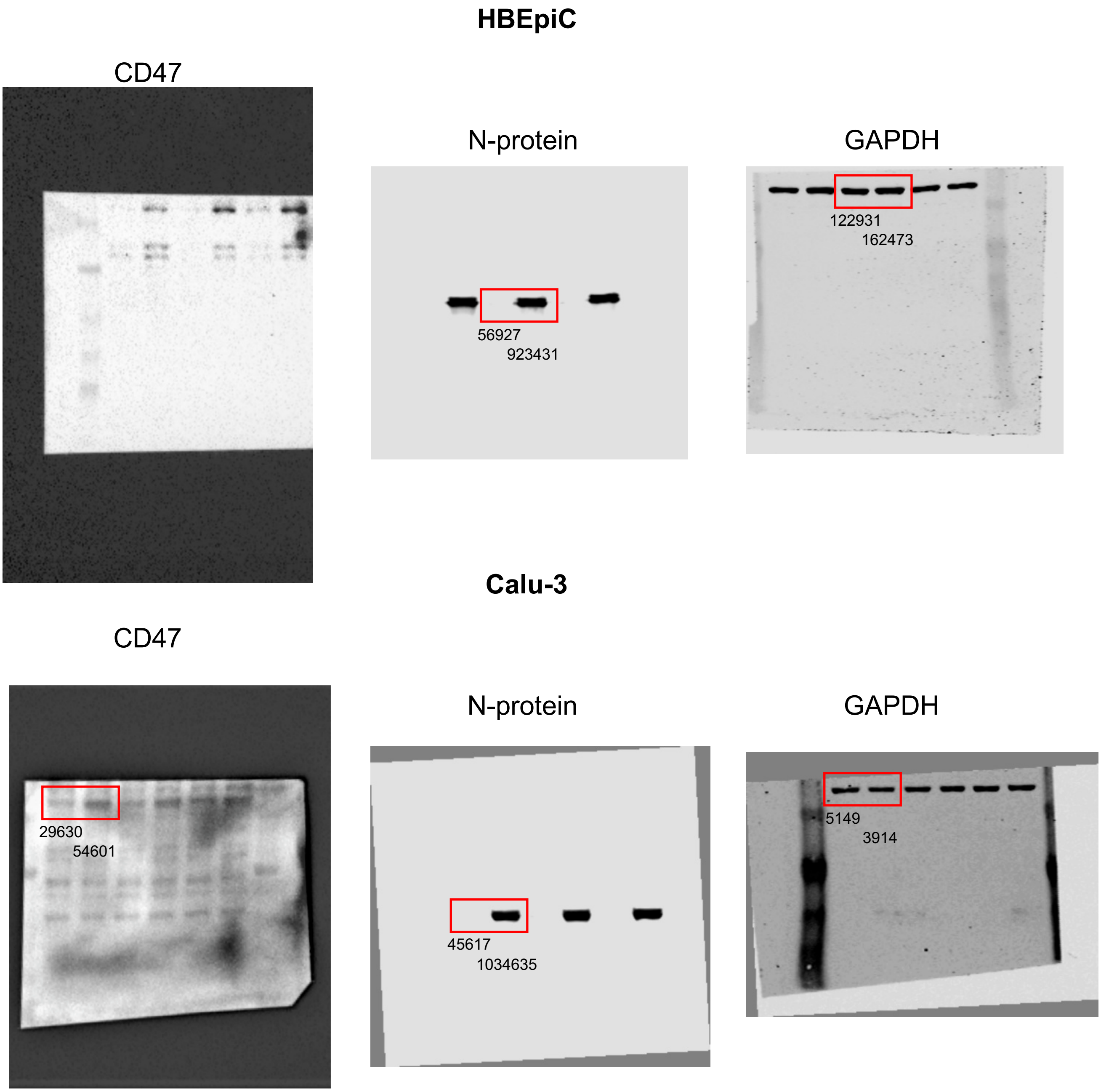


**Suppl. Figure 1**. Uncropped Western blots to Figure 1. Bands are indicated by frames. Numbers indicate quantification results.
