## Supplementary material for "CD47 as a potential biomarker for the early diagnosis of severe COVID-19": Suppl Figure 2

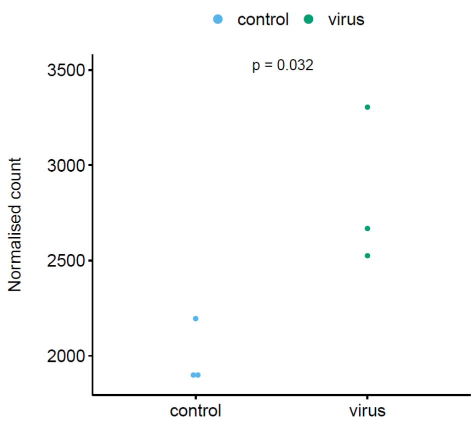


CD47

**Suppl. Figure 2**. CD47 mRNA levels in SARS-CoV-2-infected Calu-3 cells (data derived from [Blanco-Melo et al., 2020]). P-values were determined by two-sided Student's t-test.
